# A Method for the Automatic Normalization and 3D Facial Mesh Extraction from Head Magnetic Resonance Imaging

**DOI:** 10.1101/2020.07.20.211748

**Authors:** Xavier Sevillano, David Ureña, Rubèn Gonzàlez, Mar Fatjó-Vilas, Raymond Salvador, Edith Pomarol-Clotet, Neus Martínez-Abadías

## Abstract

The analysis of 3D facial shape in medicine is motivated by the fact that certain diseases and syndromes are associated to specific facial dysmorphologies. In this context, 3D facial shape analysis constitutes a promising and non-invasive support to traditional diagnostic methods. In this work, we explore the use of head magnetic resonances to obtain accurate 3D facial meshes that enable subsequent facial shape analysis. We present a fully automatic method that normalizes the orientation and alignment of 3D point clouds corresponding to head magnetic resonances by detecting salient facial features. Moreover, using clustering techniques, our method also allows to eliminate noise and artifacts appearing in magnetic resonance imaging. Finally, through bidirectional ray tracing, we obtain a dense 3D facial mesh that accurately captures facial shape. The proposed method has been built and evaluated on a dataset of 185 head magnetic resonances, and it has demonstrated its ability to successfully orient, align and obtain a dense 3D facial mesh with a high accuracy rate.

## 1 Introduction

Accumulating evidence suggests that 3D facial shape analysis can be used as a clinical biomarker in genetic disorders that present with specific facial dysmorphologies [4, 5].

Examples of this include early diagnosis of fetal alcohol spectrum disorder [12], epilepsy [1], Down syndrome [11] or schizophrenia [6].

In this context, 3D facial shape analysis constitutes a promising and non-invasive support to traditional diagnostic methods [3]. Often, this analysis is based on the detection of facial landmarks at distinctive facial features (eyes, nose, mouth) and the computation of facial biomarkers based on statistical shape analysis tools such as geometric morphometrics [8].

Typical means to obtain 3D facial data include *i)* photogrammetry systems, which combine the images captured simultaneously by several cameras to reconstruct a facial scan in the form of a 3D mesh [7], *ii)* structured light imaging, which uses the distortion suffered by a light pattern when it falls upon on a 3D object to infer the surface shape of the latter [9], or *iii)* line laser scanners, which use triangulation theory to obtain feature data of the surface from a laser sweep of the facial surface [15].

Albeit less common, magnetic resonance imaging (MRI) can be used as a source of 3D anatomical data [2]. Indeed, under appropriate scanning conditions, the resulting facial reconstructions are as accurate as 3D surface laser scans or 3D photographs [10].

This work describes an automatic method for obtaining 3D facial meshes from head magnetic resonances (HMR). The proposed method is capable of orienting, aligning and denosing, in a fully unsupervised manner, 3D point clouds corresponding to HMR. Then, it applies bidirectional ray tracing to obtain a dense mesh of the facial surface, which ultimately captures face shape, discarding unnecessary data points of the HMR. Thus, in scenarios where large volumes of data must be processed, our proposal gives an automatic alternative to cumbersome and error-prone manual processing, while saving both labor and time.

## 2 Materials and methods

### 2.1 Data collection and pre-processing

The proposed method was developed and evaluated using 185 HMR obtained at the Hospital Sant Joan de Déu in a 1.5T GE Sigma scanner. High-resolution structural 3D T1 MRI were obtained with the following acquisition parameters: matrix size 512 × 512; 180 contiguous axial slices; voxel resolution 0.47 × 0.47 × 1 mm^3^; echo (TE), repetition (TR) and inversion (TI) times, (TE/TR/TI)=3.93ms/ 2000ms/ 710ms, respectively; flip angle 15.

The stacked sets of images were loaded to the 3D analysis software for sci- entific data Amira (http://www.fei.com/software/amira-3d-for-life-sciences/) to manually reconstruct the 3D surfaces of the anatomy of the participants. Since the grayscale value of each voxel in MRI is correlated with the density of the tissue, a threshold was manually adjusted for each participant to optimize skin segmentation and eliminate the glasses they wore during the scan due to the MRI protocol, while preserving their facial anatomy. However, even after this manual process noisy residuals are still present in the HMR.

Finally, the thresholded HMR were stored as PLY files containing approximately 1 million (*x, y, z*)-coordinates points each. These constitute a discrete and highly dense representation of a human head including its internal organs.

In the data set, the HMR appear arbitrarily oriented towards the *Z*^+^ axis or the *Y* ^+^ axis. And among the latter, around half of the HMR have the top of the head pointing towards the *Z*^+^ axis, and the other half, towards the *Z*^*–*^ axis. Figure 1 presents three examples of these different orientations, and Table 1 shows the number of instances in the data set with each particular orientation.

**Table 1:** Number of instances in the data set according to their orientation.

| Orientation | Number of instances |
| --- | --- |
| $Y^+$ (top of head to $Z^+$) | 35 |
| $Y^+$ (top of head to $Z^-$) | 65 |
| $Z^+$ | 85 |

**Fig. 1:**
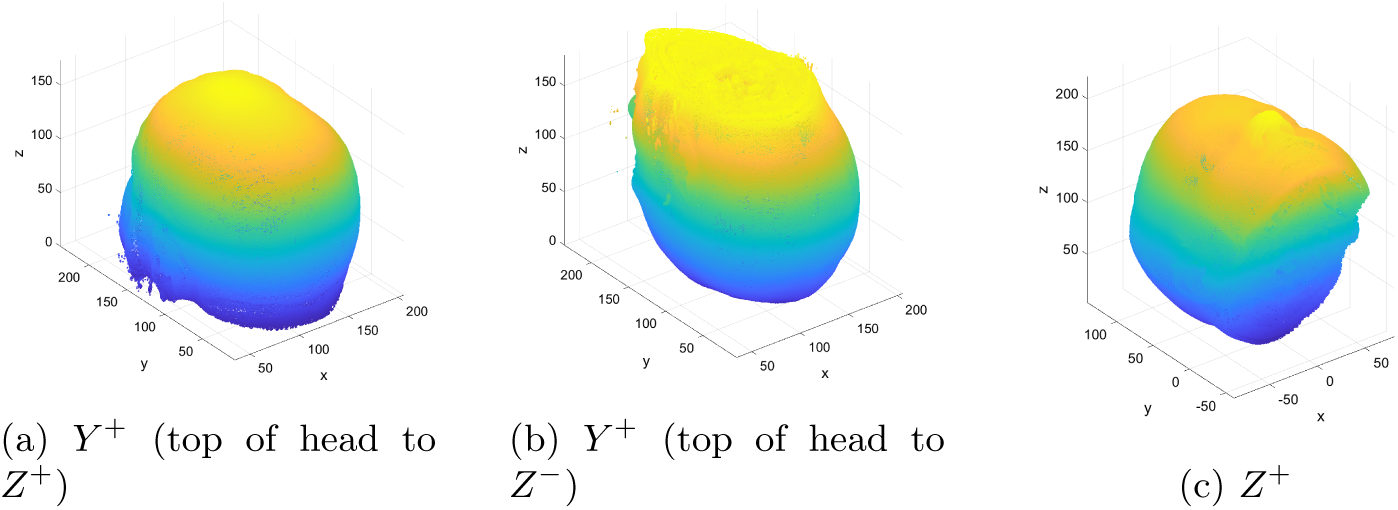
Examples of the three HMR orientations found in the data set.

### 2.2 Orientation normalization

Our method first orients the faces of the HMR automatically towards the *Y* ^+^ axis with the top of the head pointing towards the *Z*^+^ axis, following these steps:

1. **Centering** The coordinates of the HMR point cloud centroid 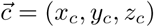 are computed and subtracted from each point, so that the point cloud center becomes the origin (0,0,0).
2. **Nose and chin detection** To automatically determine face orientation, points located in the nose and chin areas are detected in an unsupervised fashion. The tips of the nose and the chin *i)* are among the farthest features from the head centroid, and *ii)* have a characteristic pointy shape. For this reason, the Euclidean distance of each point to the origin is computed, and the 5% of the points with greater distance are considered to form point set *𝒫*_*far*_, which always contains points located near the chin and/or nose. Depending on the subject head anatomy, *𝒫*_*far*_ may also contain points located near the crown of the head. To discriminate these points, the *peakness score* (PS) of each point *p* = (*x*_*p*_, *y*_*p*_, *z*_*p*_) ∈ *𝒫*_*far*_ is computed as a local estimator of curvature. PS(*p*) is defined as the ratio of neighbours of *p* (within a pre-specified radius) that are located closer to the origin than *p*. Thus, if *p* is a pointy feature, PS(*p*) will tend to 1, while it will tend to 0.5 if it is located on a flat area. Next, points are ranked according to their PS, and only the highest ranked 15% subset of *𝒫*_*far*_ are included in the new point set *𝒫*_*far*+*peak*_. In most cases, using PS is enough to remove points near the crown of the head from set *𝒫*_*far*+*peak*_. In cases where this is not sufficient, we apply clustering [14] to unequivocally detect nose and/or chin points: the points in *𝒫*_*far*+*peak*_ are grouped according to their coordinates to discover spatial clusters with a minimum Euclidean distance of 20 between points from different clusters. This results in one or two clusters of points. If only one cluster is found, it contains points close to the nose or chin tips. When two clusters are obtained, we discern which one corresponds to the facial region by fitting spheres to the points in each cluster using the M-estimator SAmple Consensus (MSAC) algorithm [13]. The sphere with the smallest radius is the one corresponding to the nose/chin, thus enabling the unequivocal detection of the facial region. The set of points resulting from this step is denoted as *𝒫*_*nose/chin*_.
3. **Orientation correction** The maximum coordinate of the points in *𝒫*_*nose/chin*_ indicates face orientation. Then, the whole HMR point cloud is rotated by applying a 3D affine geometric transformation that depends on the initial face orientation:
  a. If the initial orientation was towards *Z*^+^, the point cloud is rotated *π* radians about the *Z* axis and then 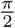 radians about the *X* axis.
  b. If the initial orientation was towards *Y* ^+^ with the top of the head towards *Z*^*–*^, the point cloud is rotated *π* radians about the *Y* axis.

### 2.3 Alignment

Once all the HMR point clouds are faced towards the *Y* ^+^ axis, the next step consists in aligning them uniformly. To this aim, the point cloud is rotated about the *X* and *Z* axes as required until the line that goes from the origin to the nose tip is perfectly aligned with the *Y* axis. The process consists in the following steps:

1. **Nose tip detection** The HMR point cloud is trimmed on its upper and lower parts (*Z* axis), to eliminate the top of the head and the mandible region, respectively. All those points whose *z* component ranges between the top 15% and the bottom 15% are included in the point set *𝒫*_*nosetip*_. Among them, the nose tip point is detected as the point with the maximum *y* component, whose coordinates are (*x*_*nose tip*_, *y*_*nose tip*_, *z*_*nose tip*_).
2. **Rotation about the** *Z* **and** *X* **axes** To align the nose tip with the *Y* axis, and given the coordinates of the nose tip point, the rotation angles about the *X* and *Z* axes are computed as:

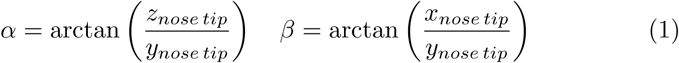

Finally, the alignment step is completed by rotating the HMR point cloud about the *X* and *Z* axes applying the *M*_*x*_ and *M*_*z*_ 3D rotation matrices:

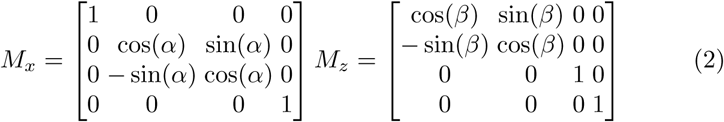

### 2.4 Denoising

The manual thresholding step described in Section 2.1, which is intended to remove the glasses worn by the patients and reconstruct their 3D facial anatomy, sometimes is unable to remove them completely. As a consequence, noisy residuals often appear in the HMR of our data set. Figure 2 presents an example of a noisy HMR.

**Fig. 2:**
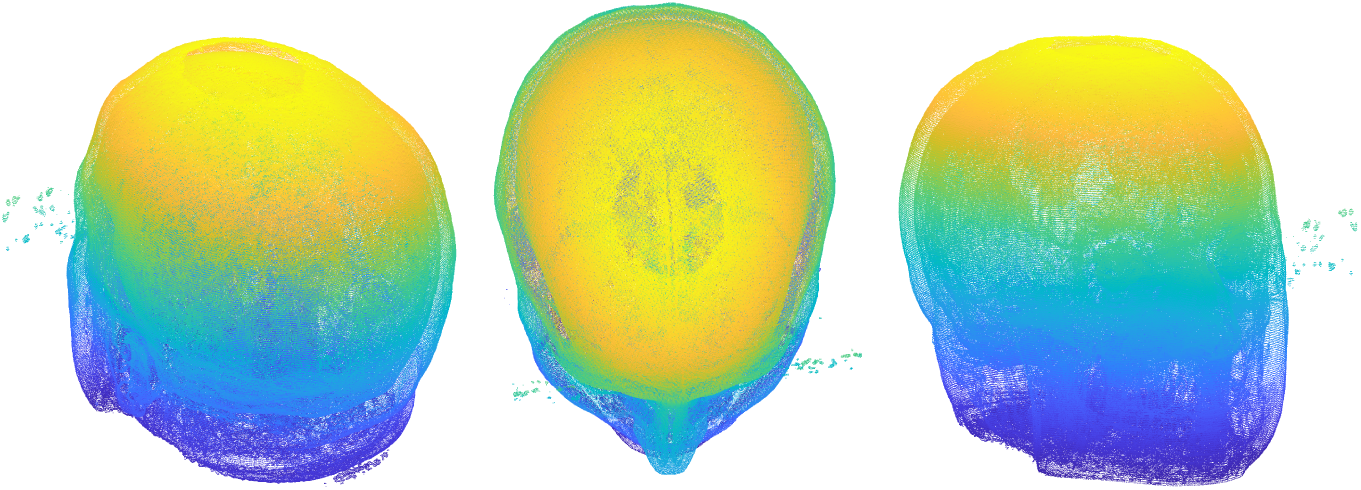
Example of noisy residuals in the HMR.

To automatically eliminate these residuals, clustering is applied to segment the point cloud. In this case, as the goal is to separate the main structure in the HMR (the head of the patient) from the noisy residuals that may surround it, the minimum Euclidean distance between points from different clusters is reduced to 1.

### 2.5 Facial mesh extraction

The HMR contain points corresponding to internal head organs. This information is irrelevant to facial shape analysis. For this reason, the last step of the method consists in extracting the 3D facial mesh from the HMR.

It consists in the following steps:

1. **Frontal region filtering** As the HMR is now oriented towards the *Y* ^+^ axis, the points corresponding to the frontal region are obtained by discarding those with a negative *Y* coordinate, thus cutting the point cloud size nearly by half.
2. **Bidirectional ray tracing** To ensure coverage of all facial areas, a dense set of uniformly distributed rays are traced from two orthogonal planes: the XZ (frontal ray tracing) and the YZ planes (lateral ray tracing). Figure 3 shows a small subset of 10 rays (drawn in green) in each direction for visualization purposes only. Each beam of parallel rays is generated by creating a uniform grid of adjustable horizontal and vertical resolution, and a ray is traced from each point of the grid. The intersection between each ray and the surface of the HMR allows to reconstruct the facial mesh. To that end, an *α*-shape is built upon the HMR point cloud, and the first point of each ray to intersect the *α*-shape is deemed as a facial mesh point.

**Fig. 3:**
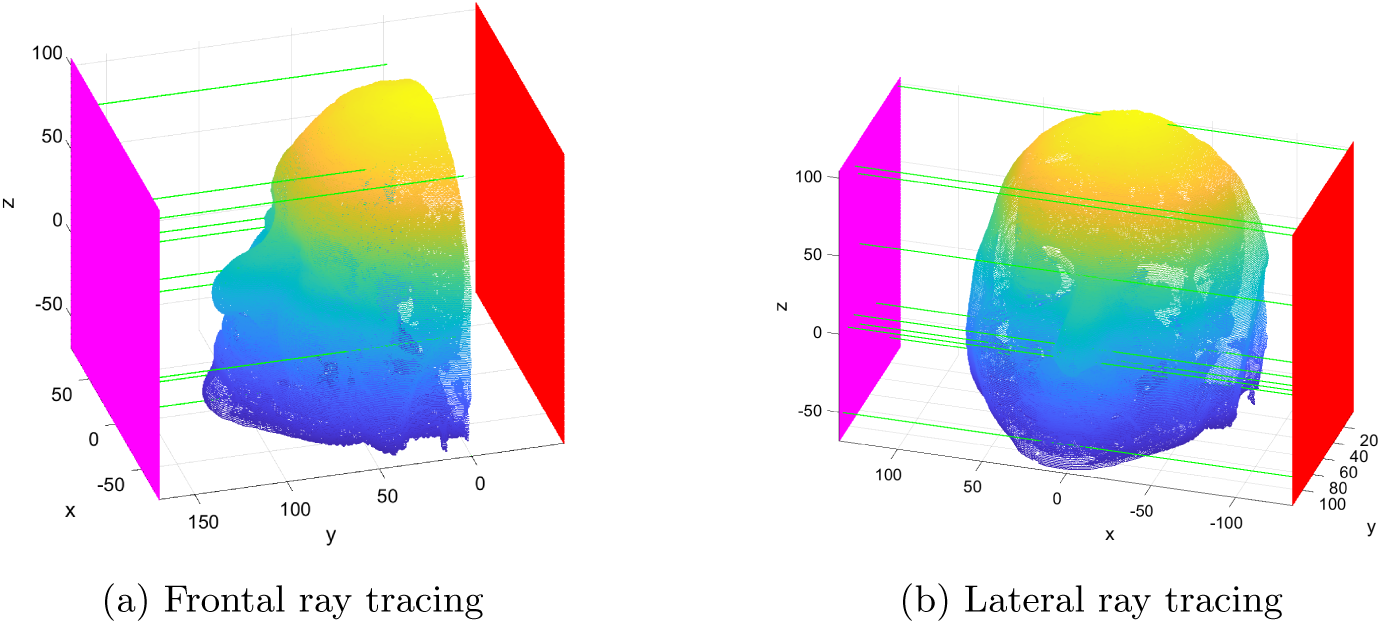
Bidirectional ray tracing.

## 3 Results

### 3.1 Orientation normalization

Figure 4 presents partial and final results of the orientation normalization process. To detect the facial region (nose and chin points), we first present two examples of the *𝒫*_*far*_ point sets, highlighted in magenta: containing only facial points (Figure 4a) and facial plus crown of the head points (Figure 4b).

Next, also highlighted in magenta, we present two examples of the *𝒫*_*far*+*peak*_ sets, containing only facial points (Figure 4c) and facial plus crown of the head points (Figure 4d).

**Fig. 4:**
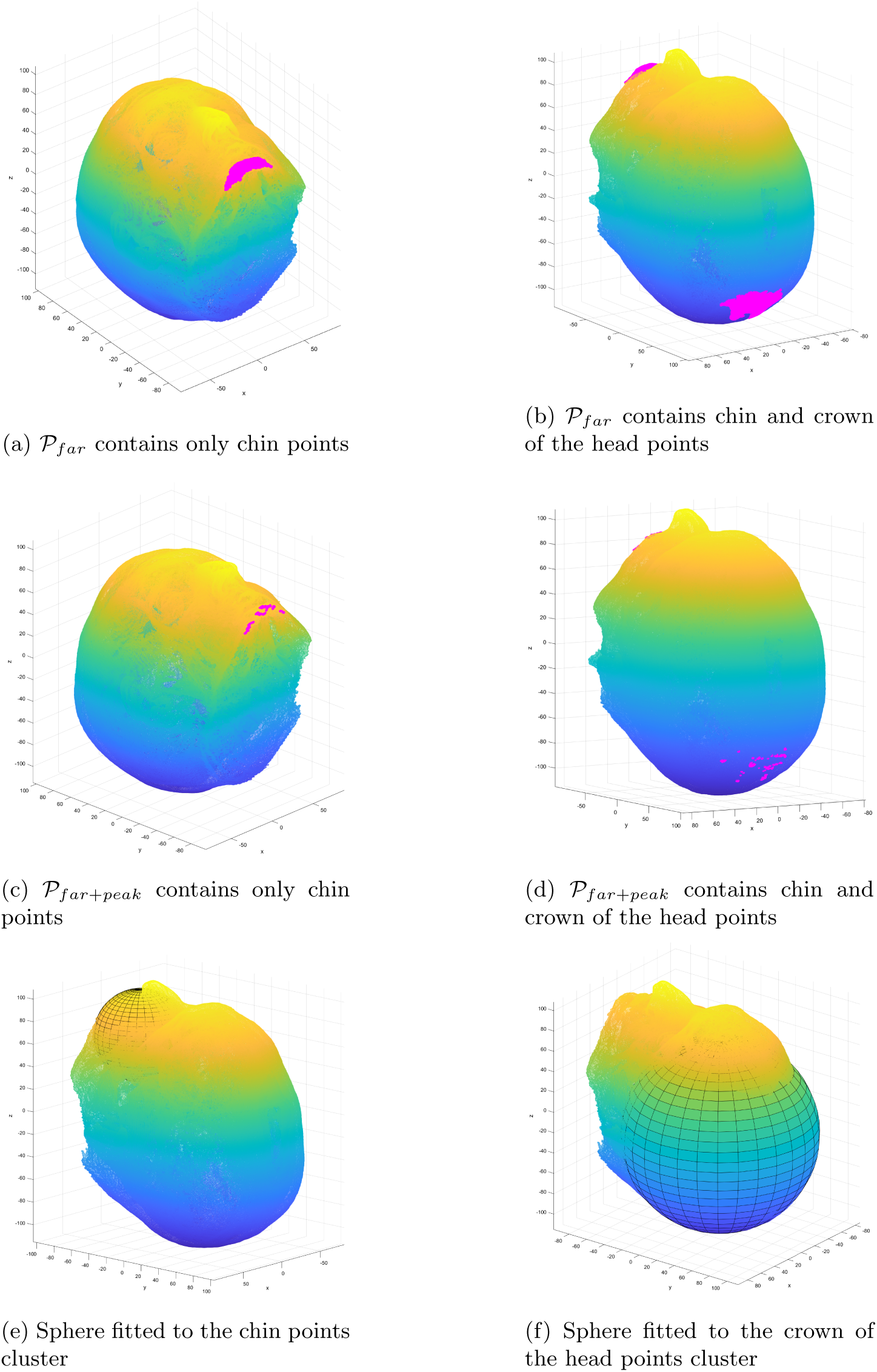
Orientation normalization results.

In this latter case, the detection of the frontal region is completed by fitting the spheres to the chin (Figure 4e) and crown of the head points (Figure 4f).

From a quantitative perspective, our method detects the facial region with a 98% accuracy, thus enabling the automatic orientation normalization of the vast majority of our data set.

### 3.2 Alignment

Figure 5 presents partial and final results of the HMR alignment process, which relies on nose tip detection.

**Fig. 5:**
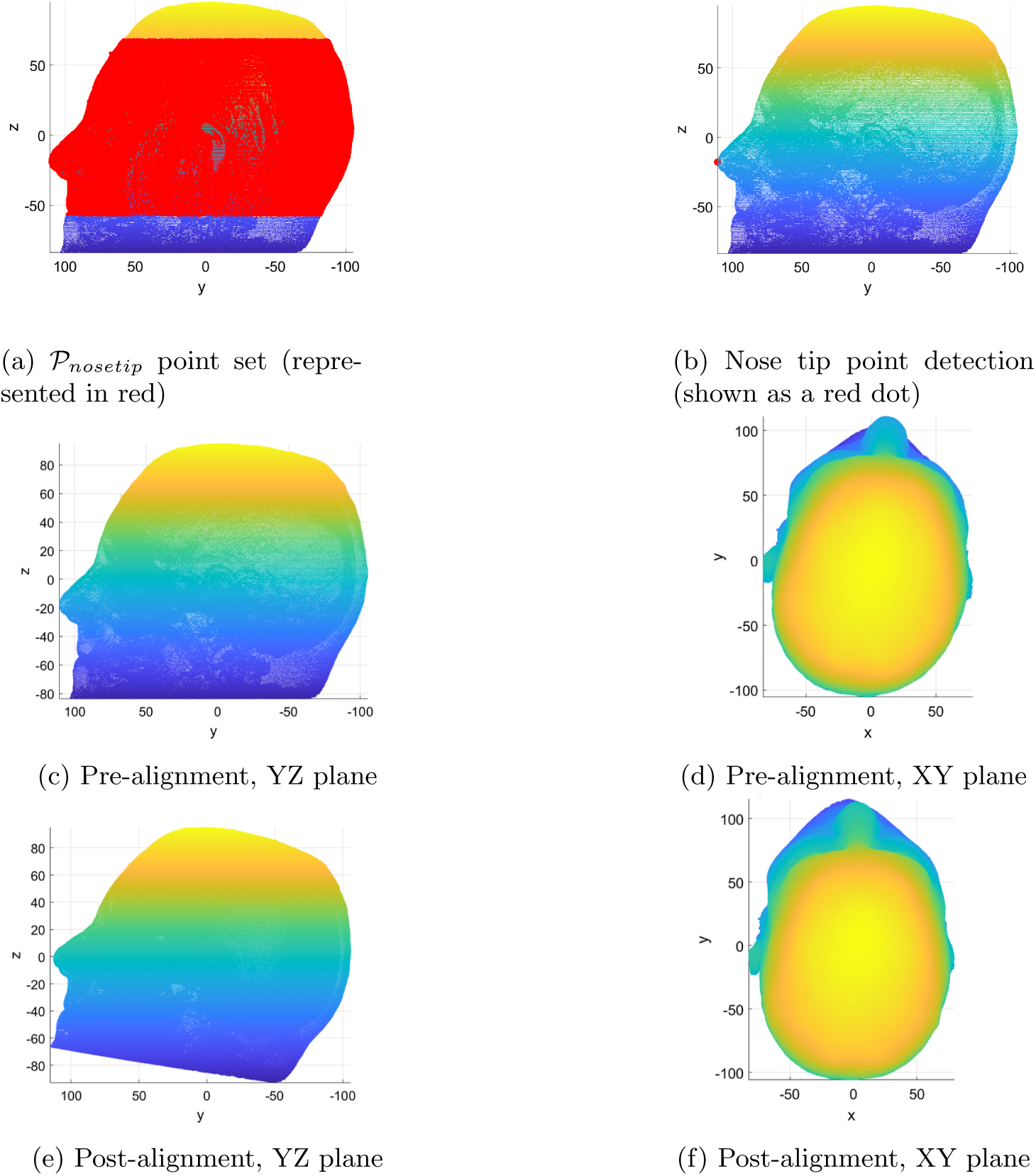
Alignment results.

First, Figure 5a depicts the point set *𝒫*_*nosetip*_ used for nose tip detection.

Next, the point in *𝒫*_*nosetip*_ with the maximum *y* component is detected as the nose tip (marked with a red dot in Figure 5b).

In quantitative terms, nose tip point is detected with a 99% accuracy, thus allowing to automatically align nearly all the HMR in our data set.

Finally, Figures 5c to 5f present the result of rotating the HMR about the X and Z axes, obtaining the aligned version of the HMR.

### 3.3 Denoising

Figure 6a depicts the clustered HMR, with each cluster represented with a dif- ferent color.

**Fig. 6:**
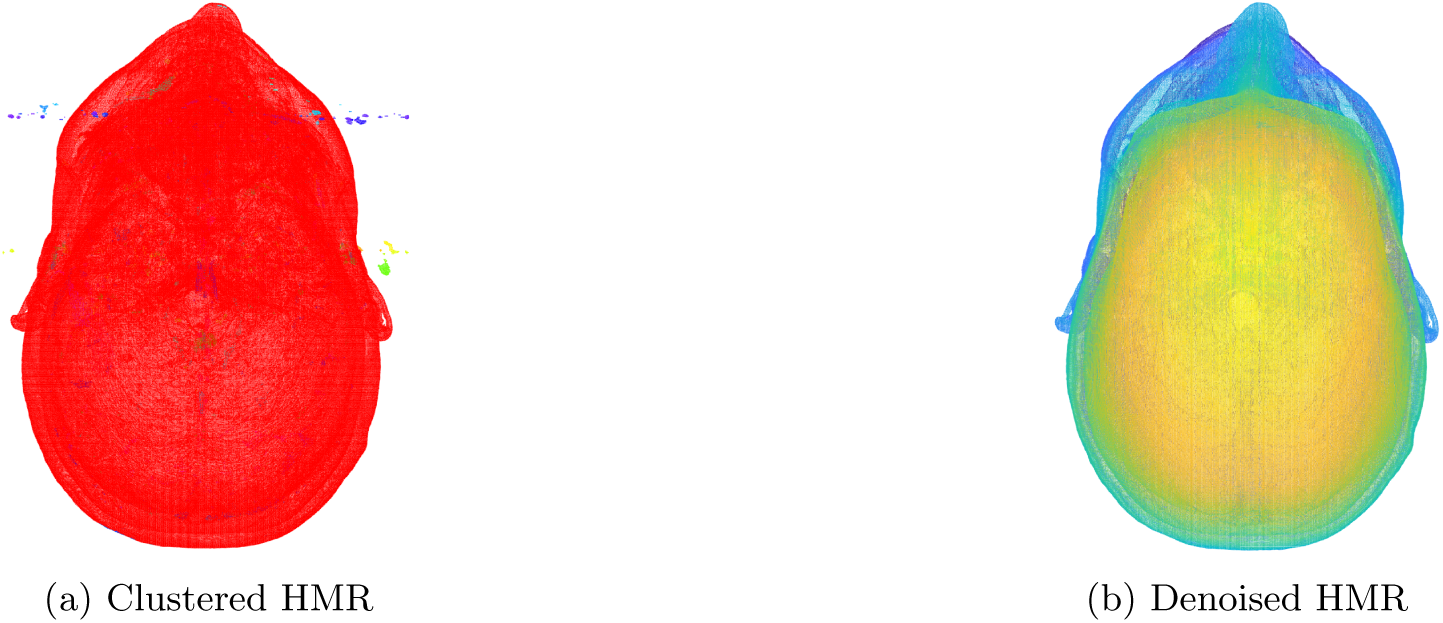
Denoising results.

Only the points assigned to the largest cluster (corresponding to the head) are kept, while the points in the less populated clusters are discarded.

Figure 6b shows an example of the results obtained by denoising via clustering.

### 3.4 Facial mesh extraction

Figure 7a depicts (in magenta) the intersection points between the dense beam of frontally traced rays and the HMR. In this case, the resolution of the ray tracing grid is 0.55, which results in over 96000 frontal rays.

**Fig. 7:**
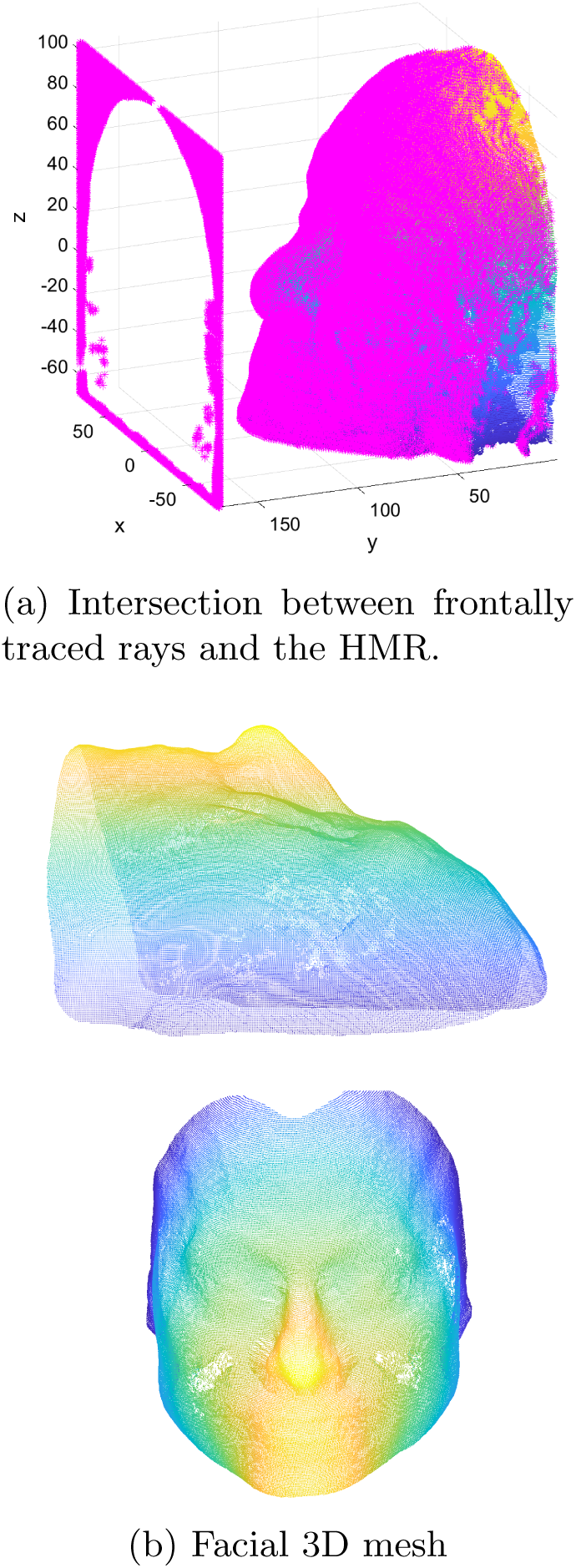
Ray tracing results.

Notice that *i)* most points are located on the face surface, *ii)* some rays do not intersect with the head due to the geometry of the head or the presence of remaining artifacts (thus, ray tracing also allows eliminating unwanted noise), and *iii)* the coverage on some areas (like nose wings) is lower, which is later compensated by the lateral ray tracing.

Figure 7b presents the final 3D facial mesh obtained after bidirectional ray tracing. Using the aforementioned resolution, a dense mesh formed by over 164000 points is obtained. It is important to notice that the structure of the most salient facial features (eyes, nose and mouth) is accurately retrieved, thus allowing the detection of landmarks for subsequent 3D facial shape analysis based on geometric morphometrics.

## 4 Conclusions

Head magnetic resonances are a source for obtaining accurate 3D facial data.

This work has introduced a fully automatic method for obtaining normalized facial 3D meshes from head magnetic resonances.

Our proposal, evaluated on a testbed of 185 head magnetic resonances, successfully normalizes the orientation and alignment of the magnetic resonance images. Moreover, the method uses bidirectional ray tracing of adjustable resolution, thus allowing to obtain dense facial 3D meshes that capture face shape accurately.

This method simplifies the use of HMR for 3D facial shape analysis that enables clinical diagnostic applications for many syndromes with associated facial dysmorphologies [3].

Our next steps are oriented towards the automatic detection of facial landmarks on the obtained 3D facial meshes, and the use of artificial intelligence for the diagnosis of diseases with associated subtle facial phenotypes.

## Notes

### Competing Interest Statement

The authors have declared no competing interest.

